# Treating Influenza and SARS-CoV-2 via mRNA-encoded Cas13a

**DOI:** 10.1101/2020.04.24.060418

**Authors:** Emmeline L. Blanchard, Daryll Vanover, Swapnil Subhash Bawage, Pooja Munnilal Tiwari, Laura Rotolo, Jared Beyersdorf, Hannah E. Peck, Nicholas C Bruno, Robert Hincapie, M.G. Finn, Frank Michel, Eric R. Lafontaine, Robert J. Hogan, Chiara Zurla, Philip J. Santangelo

## Abstract

Here, Cas13a has been used to target and mitigate influenza virus A (IAV) and SARS-CoV-2 using a synthetic mRNA-based platform. CRISPR RNAs (crRNA) against PB1 and highly conserved regions of PB2 were screened in conjunction with mRNA-encoded Cas13a. Screens were designed such that only guides that decreased influenza RNA levels in a Cas13-mediated fashion, were valid. Cas13a mRNA and validated guides, delivered post-infection, simulating treatment, were tested in combination and across multiplicities of infection. Their function was also characterized over time. Similar screens were performed for guides against SARS-CoV-2, yielding multiple guides that significantly impacted cytopathic effect. Last, the approach was utilized *in vivo*, demonstrating the ability to degrade influenza RNA in a mouse model of infection, using polymer-formulated, nebulizer-based mRNA delivery. Our findings demonstrate the applicability of Cas13a in mitigating respiratory infections both *in vitro* and in a mouse model, paving the way for future therapeutic use.

## Introduction

There are 219 species of viruses that are known to infect humans ^1^ of which, 214 are RNA viruses ^2^. It is estimated that viral infections contribute to approximately 6.6% of global mortality ^3^. This is especially concerning, given that there are approximately 90 drugs (since 1963-2016) to treat only 9 viral species ^4^. In addition, there are approved vaccines for only 15 viral species. Reassortment, antigenic shift and drift, for influenza, as well as antibody dependent enhancement (ADE) for SARS and possibly, SARS-CoV-2, pose challenges to vaccine development^5 6^. These factors likely contribute to epidemics and pandemics. Human health is thus under constant threat due to emerging and reemerging viral infections ^7^. Outbreaks of Zika ^8^, Ebola ^9^, and the current SARS-CoV-2 pandemic^10^, and the potential for future influenza pandemics^11^, warrant the development of new classes of anti-viral drugs^4^. Current drug development is focused on small molecules and neutralizing antibodies, which require high doses or frequent re-dosing to obtain functional outcomes ^12,13^. Thus, it is crucial to address the need for antivirals that are broad spectrum, flexible and effective across multiple viral species or strains.

The discovery of a RNA targeting, class II - type VI CRISPR-Cas system in bacteria has engendered tremendous interest for potential applications. The Cas13a: crRNA complex activates when the target RNA (trRNA) complements with crRNA, and the Cas protein initiates RNA cleavage^14,15^, due to the higher eukaryote and prokaryote nucleotide-binding (HEPN) domain ^16^. This property has been used to detect specific transcripts in mixtures of nucleic acids ^15^. The specific RNase activity of Cas13a and Cas13d ^17^ was also used to knockdown endogenous genes, and more recently, two groups have demonstrated the ability of Cas13b or Cas13d to degrade influenza RNA^18,19^. In both cases, transient transfection with plasmids or stable cell lines expressing Cas13 were used prophylactically against influenza RNA. In each case, Cas13 was expressed at least 24 hrs prior to infection. This prior work was clearly useful to demonstrate proof of principle.

Here we examined crucial steps towards using this approach as a treatment for respiratory viral infections. Given the advantages of transient expression for treating infections, we developed synthetic mRNA to express LbuCas13a both with and without an NLS sequence, as influenza RNA can be localized to both compartments at different times post-infection, and both constructs were tested for function within in-tube assays. Why synthetic mRNA? Synthetic mRNA expression is transient, has very little chance of integrating, and innate immune responses can be mitigated through sequence design, modified nucleotides, and purification ^20–22^. In addition, given Cas13a is a bacterial protein, it is possible there may be immune responses to the protein itself and this is more likely if it is expressed for long periods of time. Transient expression with minimal innate responses (no adjuvant effect) gives us the highest probability of repeat dosing *in vivo*. Thus, this approach may be crucial for repeat dosing of Cas13a in infected individuals without immune clearance of treated cells due to Cas13a mediated immune responses.

The mRNA-based approach was initially benchmarked by examining knockdown of endogenous genes, using previously published target regions^10^. In examining these genes, and expanding the number of controls used, it was found that not all knockdown of RNA was mediated by Cas13a. A number of controls was needed to ensure that the knockdown was Cas13a mediated, and that enzymatic action was occurring. Once it was determined which controls were important, guides against PB1 were screened via qPCR, given post-infection, simulating a treatment strategy. Why polymerase genes? They are more highly conserved, and without polymerase protein neither viral mRNA nor replication can occur ^10,23,24^. A potent guide against PB1 mRNA was found using this approach but guides with broader sequence homology across flu strains were sought. By first examining homology among 108 vaccine strains, a stretch of nucleotides in PB2 were found to overlap most H1N1 strains, the H1N1 pandemic strain from 2009, and H3N2 strains used in prior vaccines. Vaccine strains are a benchmark for circulating strain variation over time and thus good candidates. At least two potent guides against this sequence were found. Subsequent tests post-infection included combining guides, and time-course studies yielding potent knockdown of influenza RNA. We then asked whether this same approach could be used against SARS-CoV-2, the virus causing the current pandemic. Here, we delivered the mRNA prophylactically, as the virus is especially potent in VeroE6 cells, and cytopathic effect (CPE) was used as the metric for success. Here we demonstrate clear mitigation of SARS-CoV-2 CPE.

Last, we demonstrated that this approach has translational potential by delivering Cas13 mRNA and guides formulated with a PBAE-based polymer ^25^ via nebulizer post-influenza infection, simulating treatment. At day 3 post infection, degradation of influenza RNA in the lung was evaluated demonstrating robust knockdown. Overall, we feel that we have demonstrated the necessity of controls for designing crRNAs, the possibility of pan-influenza targeting of viral strains, and the ability to target relevant and emergent respiratory pathogens, both *in vitro* and *in vivo*.

## RESULTS

### mRNA expressed Cas13a cleaves RNA in the presence of both crRNA and target RNA

Using the rabbit reticulocyte lysate, Cas13a and Cas13a-NLS (version with nuclear localization sequence) mRNAs were translated *in vitro* and used to assess the RNA cleavage activity of the expressed Cas13a protein in conjunction with IAV crRNAs and target RNAs (trRNAs) (**Fig. 1a and Supplementary Table 1**). A variant including an NLS sequence was included for testing because influenza virus replicates in the nucleus. crRNA and trRNA were derived from a genome segment of IAV. RNaseAlert™ substrate fluorescence was the output of RNA cleavage. Cas13a and Cas13a-NLS RNA cleavage generated fluorescence increased to its maximum during the initial 10- and 20-min period, respectively, and then gradually decreased over time, likely due to photobleaching. The overall trend of RNA cleavage was similar for both the Cas13a and Cas13a-NLS (**Fig. 1b**). Our results indicate that *in vitro* translated Cas13a mediated RNA cleavage is specific and occurred only when both the crRNA and trRNA were present. In addition, when a non-target crRNA (NTCR) was used along with the trRNA, no RNA cleavage was observed, thus demonstrating crRNA-specific cleavage of trRNA by Cas13a (**Fig, 1b**). Overall, the mRNAs designed express functional Cas13 proteins.

**Fig 1.**
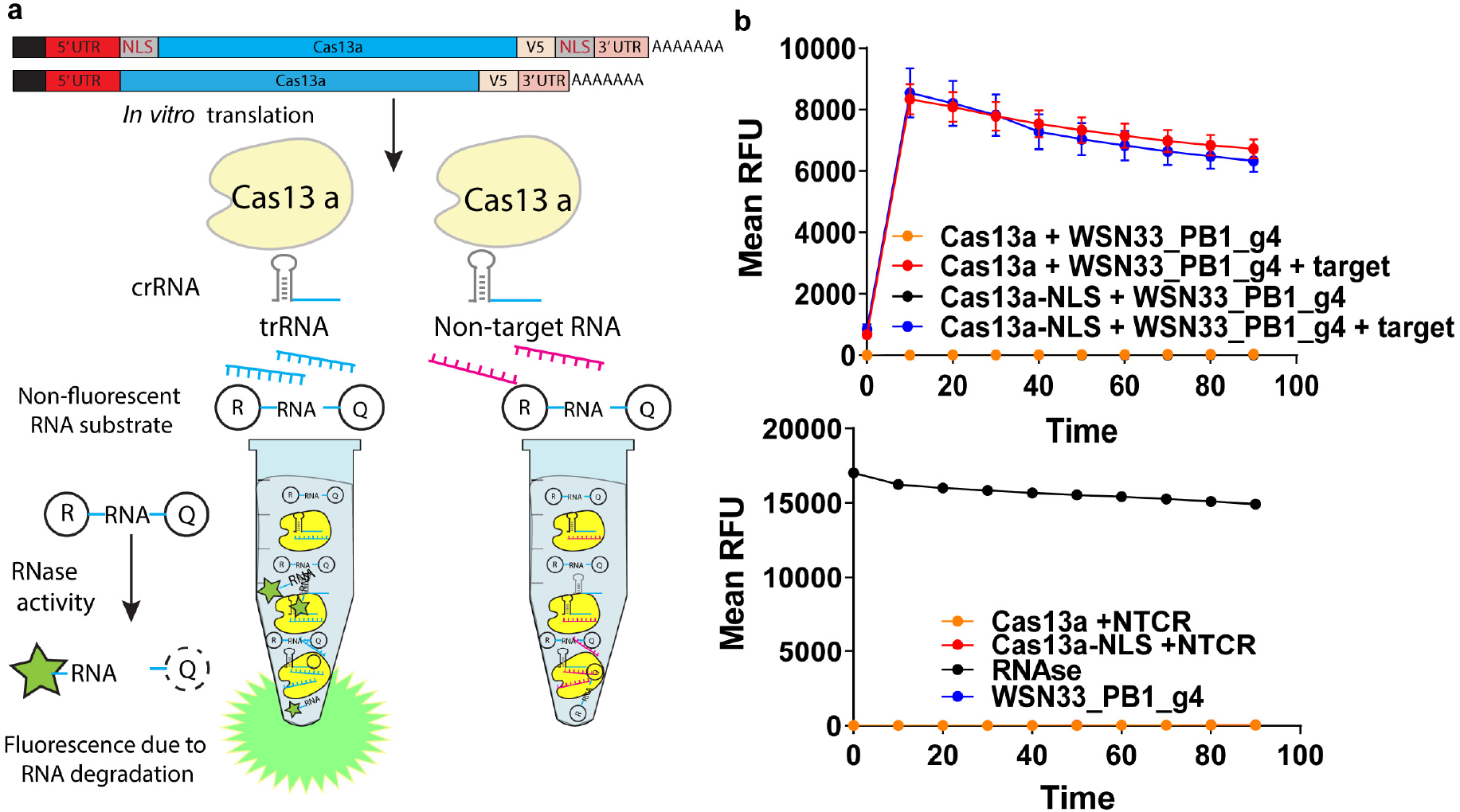
Cas13a IVT mRNA generates functional Cas13a protein. **a)** Schematic representation of the *in vitro* RNA cleavage assay. **b)** Cas13a-V5 and Cas13a-V5-NLS mRNA were translated using a rabbit reticulocyte lysate system. The proteins were mixed with RNAse Alert substrate, a guide RNA and a target RNA and the resulting fluorescence was measured over time. Means and standard error of means are shown, where n = 3 technical replicates. Negative controls include no target RNA, a non-targeting guide RNA (NTCR), and guide alone with no Cas13a protein. Positive control is RNAse.

### Endogenous gene knockdown via mRNA-based Cas13 expression

In order to more fully validate the mRNA-based approach, mRNA expressing LbuCas13a, with and without an NLS, and guides targeting PPIB, CXCR4 and KRAS endogenous genes were tested, and gene knockdown evaluated at 24 hours via qPCR (**Fig. 2a-c and Supplementary Table 2**). Initially, the knockdown relative to the non-targeted control RNA (NTCR) was evaluated, but the effects of transfection, etc., on gene expression were not being assessed. To date, most studies only evaluate targeted reduction of RNA relative to the reduction due to a NTCR. Two additional controls were added, a mRNA expressing a “dead” or inactive Cas13a plus targeted guide, and a GFP encoding mRNA plus targeted guide. Each control yielded valuable information regarding how the guide was performing. The dead version gives insight into whether “binding-only” events may affect RNA levels, while the GFP control adds additional information regarding whether the guide alone has knockdown effects. If all the controls do not affect RNA levels and knockdown is only due to the active enzyme, then enzymatic action is driving the reduction. There are cases where the Cas13a inactive version may also decrease RNA levels (**Fig. 2 b**). This is not necessarily a problem *per se*, but this control group helps distinguish “binding” related knockdown from enzymatic effects. All this information is important for establishing the mode of action of this therapeutic approach.

**Fig 2.**
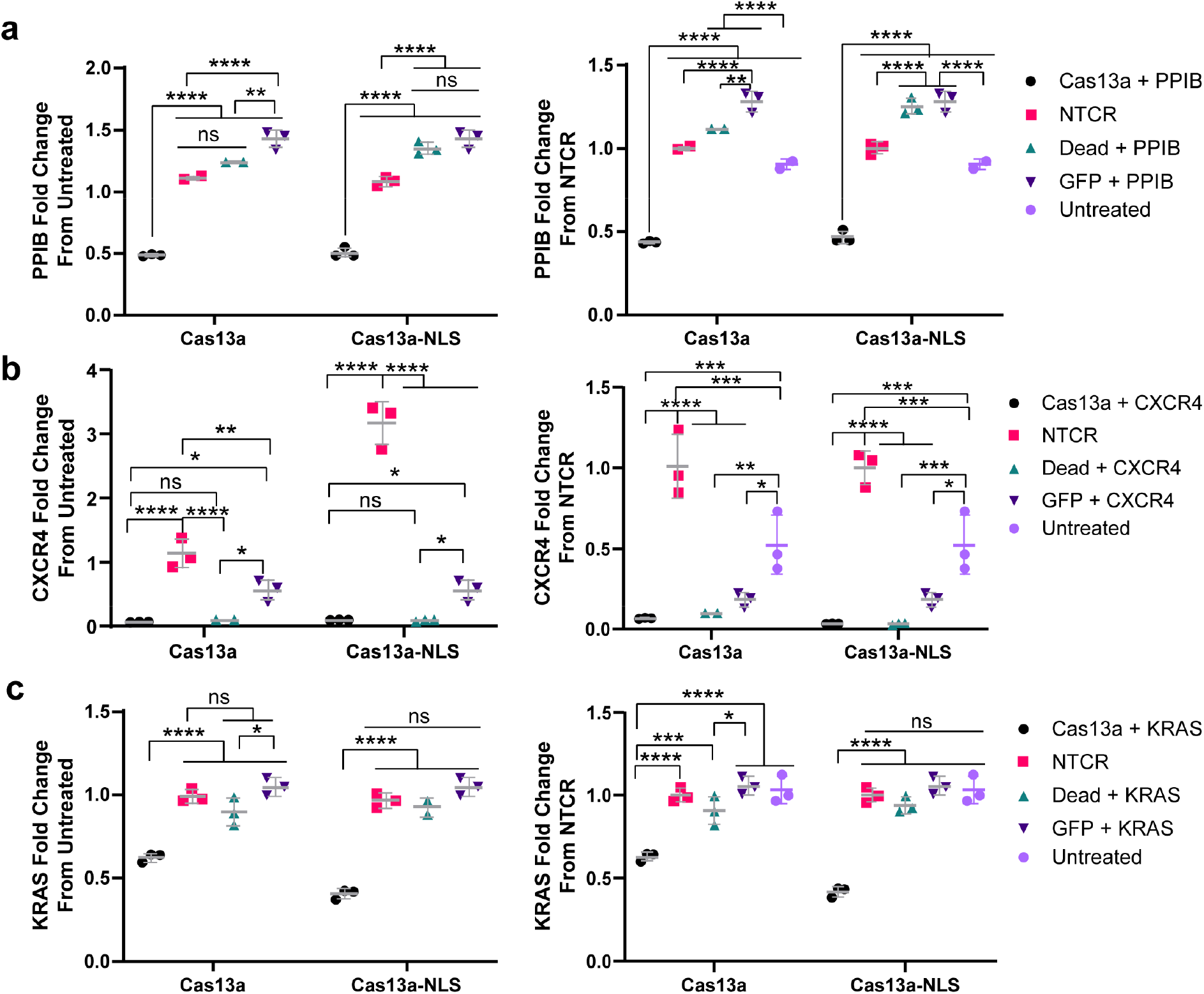
IVT Cas13a mRNA reduces endogenous mRNA levels. A549 cells were transfected with 0.5 μg of Cas13a – V5 or Cas13a-V5-NLS, dCas13a-V5, dCas13a-V5-NLS mRNA or an equal molar amount of GFP mRNA (0.2 μg) and with a 1:20 mol ratio crRNA or NTCR guides via Messenger Max. 24 h post transfection, cells were lysed and RNA was extracted. RT-qPCR was performed using GAPDH as control. Means and standard deviations are shown in grey, with n = 3 biological replicates. RT-qPCR technical triplicates were performed to determine biological replicate values. **a)** Cas13a mediated PPIB knockdown. Two-ANOVAs with Tukey multiple comparisons were performed, where ** p < 0.0035, and **** p < 0.0001. **b)** Cas13a mediated CXCR4 knockdown. Two-ANOVAs with Tukey multiple comparisons were performed, where * p <0.033, ** p < 0.005, *** p < 0.00081, and **** p < 0.0001. **c)** Cas13a mediated KRAS knockdown. Two-ANOVAs with Tukey multiple comparisons were performed, where * p < 0.04, *** p = 0.0001, and **** p < 0.0001.

For PPIB, the controls in general acted as expected, though there was some effect of the NTCR in the NLS case. The PPIB guides exhibited 57% and 53% knockdown via the cytosolic and NLS versions of LbuCas13a in 24 hrs, respectively (**Fig. 2a**). For CXCR4, unfortunately, the guides themselves, using published spacer sequences, without Cas13a expression, significantly knocked down the mRNA by over 90% (**Fig. 2b**). For KRAS, this did not occur, and instead the guides exhibited Cas13 mediated knockdown, 38% and 60%, via the cytosolic and NLS versions of LbuCas13a in 24 hrs, respectively (**Fig. 2c**). The data resulting from the CXCR4 experiments were surprising, but clearly demonstrated the need for the additional controls when screening guides against specific RNAs to ensure Cas13 mediated knockdown.

### Screening of crRNAs targeting PB1 post-infection

Next, six PB1 guides targeting both the genome and mRNA were designed and screened post-infection **(Supplementary Table 2).** Here, A549 cells were first infected with influenza A/WSN/33 at a MOI of 0.01; 24 hours post-infection, mRNA and guides were delivered, and at 48 hrs post-infection, PB1 RNA levels were evaluated by qPCR. From this initial screen 1 mRNA targeted guide (WSN33_PB1_m5) exhibited very strong knockdown, approximately 83% using the cytosolic Cas13a and 78% via the NLS version, during the 24 hr period (**Fig. 3a, Supplementary Fig.1**). None of the controls demonstrated “guide-only” effects (**Fig. 3b, c and Supplementary Fig.1**) and even the dead version showed minimal knockdown, demonstrating that the effects were likely only from enzymatic action.

**Fig 3.**
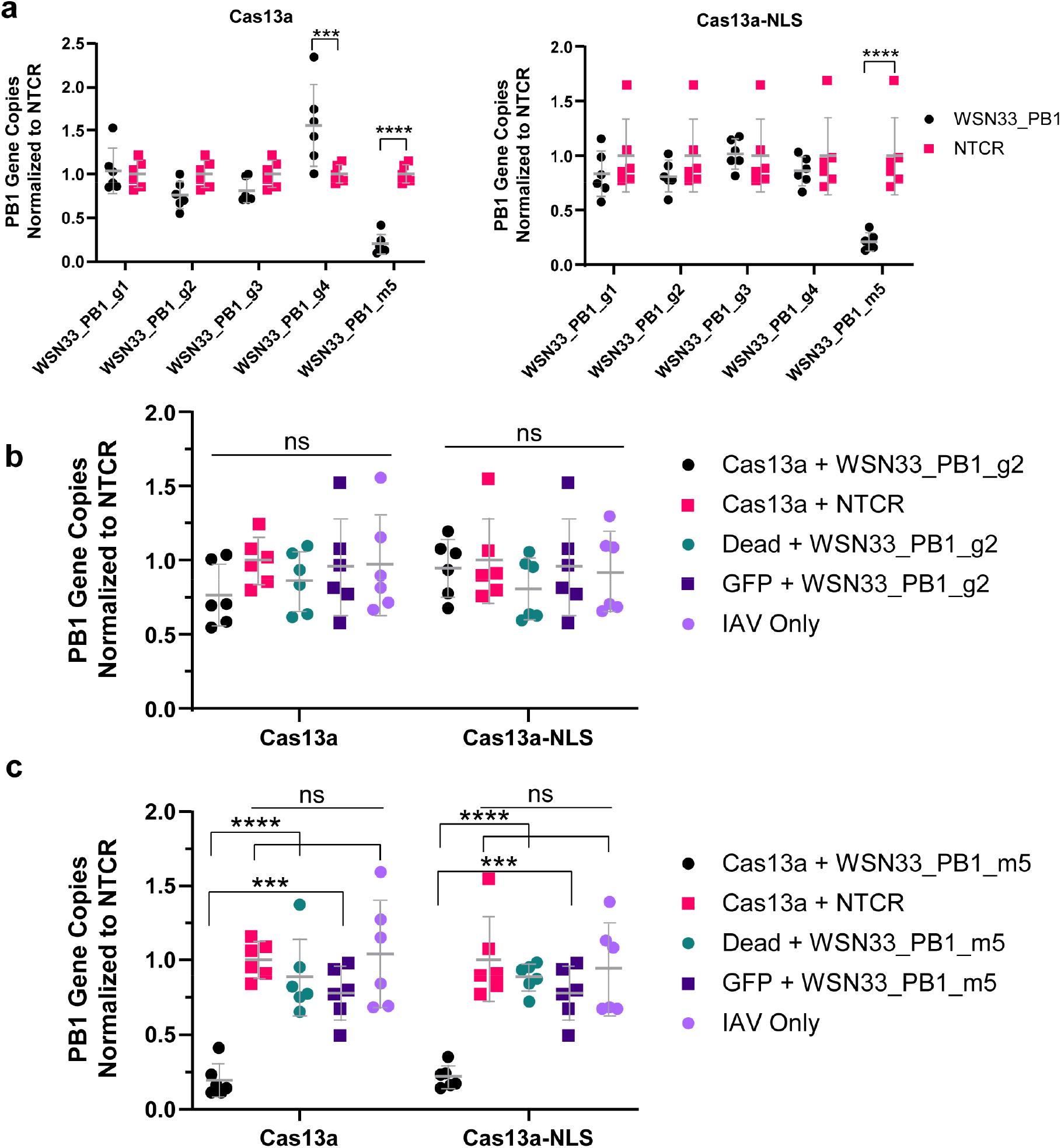
Cas13a-mediated IAV PB1 RNA knock down post-infection. **a)** A549 cells were infected with IAV A/WSN/33 at MOI 0.01. 24 hpi, cells were transfected with 0.5 μg of Cas13a-V5 or Cas13a-V5-NLS and 1:50 mol ratio of WSN33_PB1 or NTCR guides via Messenger Max. Two-ANOVAs with Sidak’s multiple comparisons were performed, where ** p = 0.0001 and **** p < 0.0001. **b)** A549 cells were infected with IAV A/WSN/33 at MOI 0.01. 24 hpi, cells were transfected with 0.5 μg Cas13a – V5 or Cas13a-V5-NLS, dCas13a-V5, dCas13a-V5-NLS mRNA or an equal molar amount of GFP mRNA (0.2 μg) with a 1:50 mol ratio of WSN33_PB1_g2 or NTCR guides via Messenger Max. A two-ANOVA with Sidak’s multiple comparisons was performed, where no significant differences were found. **c)** A549 cells were infected with IAV A/WSN/33 at MOI 0.01. 24 hpi, cells were transfected with 0.5 μg Cas13a – V5 or Cas13a-V5-NLS, dCas13a-V5, dCas13a-V5-NLS mRNA or an equal molar amount of GFP mRNA (0.2 μg) with a 1:50 mol ratio of WSN33_PB1_m5 or NTCR guides via Messenger Max. A two-ANOVA with Sidak’s multiple comparisons was performed, where *** p < 0.00065, and **** p < 0.0001. In all experimental conditions, cells were lysed and RNA was extracted 24 hpt (48 hpi). RT-qPCR was performed for WSN/33_PB1 by absolute copy number quantification. Means and standard deviations are shown in grey, with n = 6 biological replicates. RT-qPCR technical triplicates were performed to determine biological replicate values. WSN/33_PB1 gene copy numbers were normalized by NTCR values.

### Expanding the breath of guides against influenza: targeting PB2

Clearly, for this approach to have high therapeutic value, the ability to target many strains of a given virus is advantageous. For influenza, a bioinformatic comparison of 108 influenza historical vaccine strains and candidate vaccine strains, with publicly available genome sequences, yielded a number of conserved regions across different subsets of strains (**Fig. 4**). We specifically focused on our so-called “Human” subset, which included human seasonal H1N1, H1N1pdm09, and human seasonal H3N2 strains. From this analysis, a conserved region in PB2 was found, and six guides were designed against both viral mRNA and the genome of influenza **(Supplementary Tables 2 and 3).** From this exercise, three guides where found to knockdown the viral RNA >50%, delivered 24 hrs post-infection and evaluated at 48 hrs post-infection, with varying effectiveness in the nucleus and cytosol (**Fig. 5 and Supplementary Fig 2**). Guides WSN33_PB2_m4 and WSN33_PB2_g2 both exhibited over 75% knockdown at the RNA level (**Fig. 5 and Supplementary Fig. 2**) These two guides were examined for breadth, and when the combination was compared with over 52,000 strains downloaded from, over the last 100 years, they were exact matches with 99.1% of strains **(Fig. 4 e,f)**. This result was very encouraging towards the goal of a pan-influenza approach.

**Fig 4.**
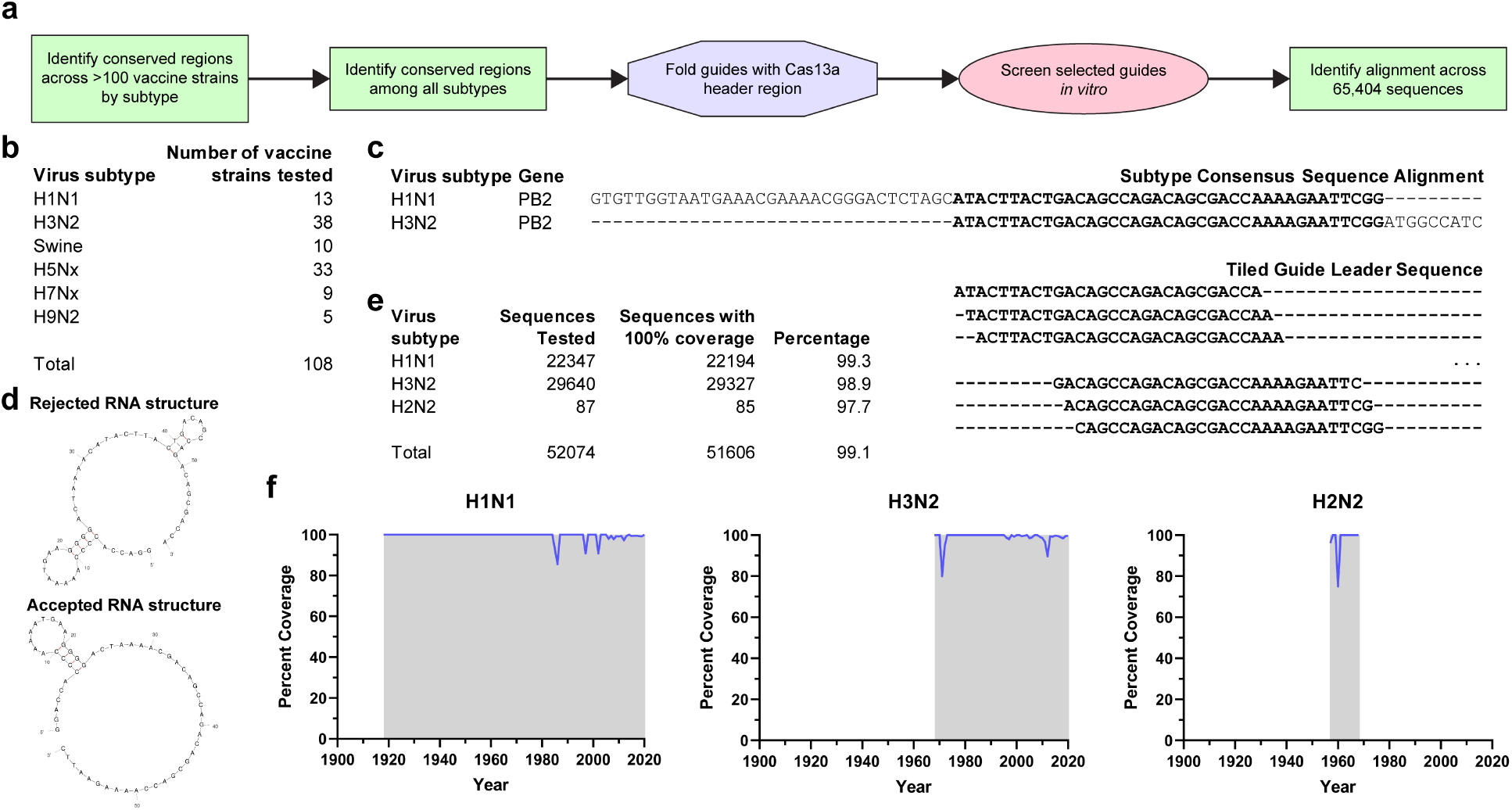
PB2 targeted guide selection and influenza A sequence coverage. **a)** Schematic overview of PB2 guide selection, validation, screening, and coverage analysis. **b)** Breakdown of influenza A vaccine candidate subtypes used for consensus searches. H1N1 – human seasonal H1N1 and H1N1pdm09 vaccine strains; H3N2 – human seasonal H3N2 vaccine strains; Swine – swine-origin H1Nx(v) and H3N2 (v) vaccine candidates; H5Nx – avian and avian-origin H5N1, H5N6, and H5N8 vaccine candidates; H7Nx – avian and avian-origin H7N9 and H7N7 vaccine candidates; H9N2 – avian and avian-origin H9N2 vaccine candidates. **c)** Consensus regions alignments between the H1N1 and H3N2 subtypes. Guides were then tiled by single-nucleotide shifts along the conserved region. **d)** Guides (with the Cas13a-Lbu header region) are folded and only accepted when no folds exist beyond the initial direct-repeat stem-loop in the header region. **e)** After *in vitro* guide screening, the top two performers, WSN33_PB2_g2 and WSN33_PB2_m4 were aligned across 52074 influenza A sequences since 1918. **f)** Percent coverage of both WSN33_PB2_g2 and WSN33_PB2_m4 across H1N1, H3N2, and H2N2 sequences. Gray region indicates years in which sequence data was available.

**Fig 5.**
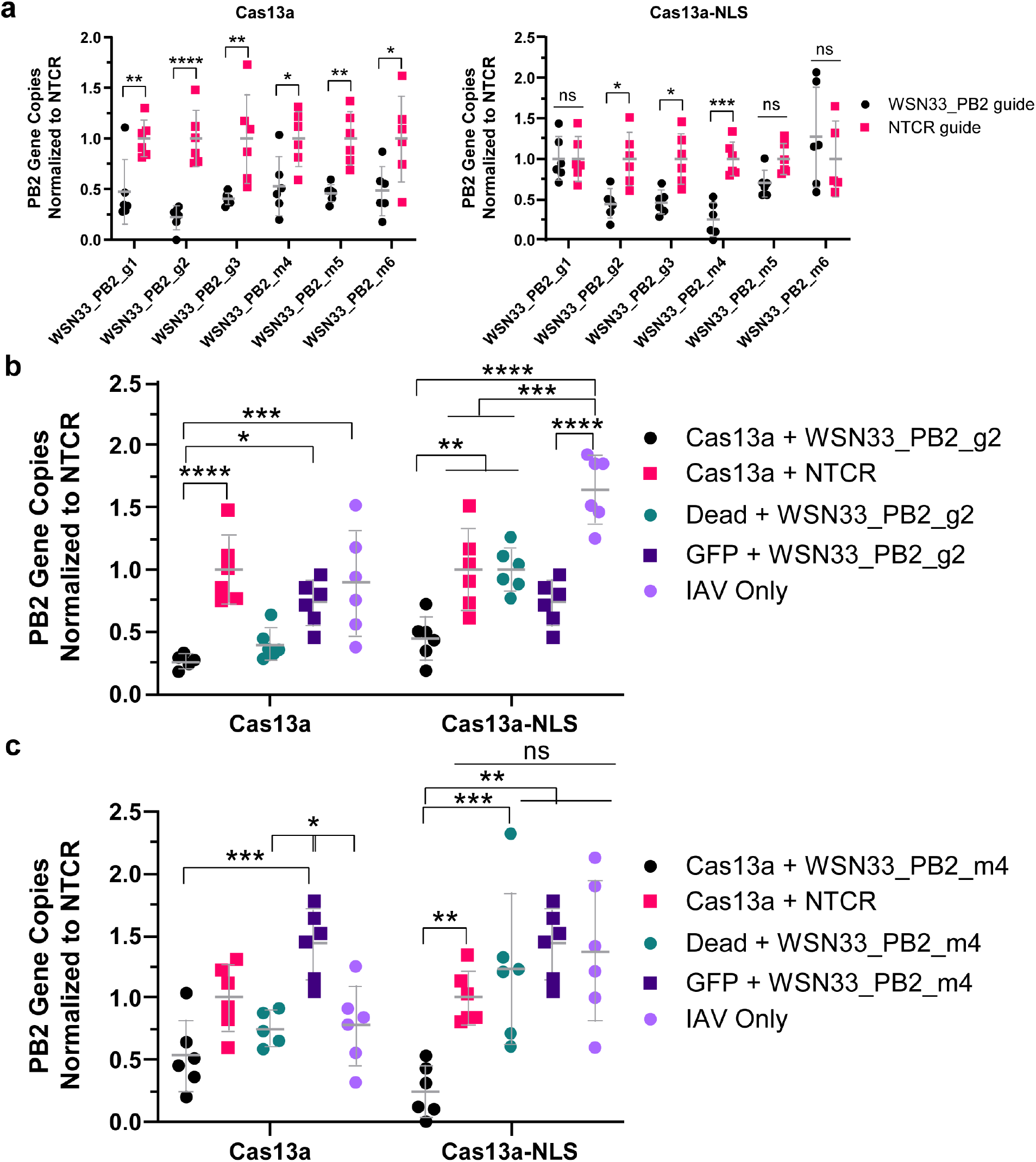
Cas13a-mediated IAV PB2 RNA knock down post-infection with broadly targeted crRNAs. **a)** A549 cells were infected with IAV A/WSN/33 at MOI 0.01 cells were transfected with 0.5 μg of Cas13a-V5 or Cas13a-V5-NLS and 1:50 mol ratio of WSN33_PB2 or NTCR guides via Messenger Max. Two-ANOVAs with Sidak’s multiple comparisons were performed, where * p < 0.026, ** p < 0.0083, *** p = 0.0005, and **** p < 0.0001. **b)** A549 cells were infected with IAV A/WSN/33 at MOI 0.01. 24hpi, cells were transfected with 0.5 μg Cas13a – V5 or Cas13a-V5-NLS, dCas13a-V5, dCas13a-V5-NLS mRNA or an equal molar amount of GFP mRNA (0.2 μg) with a 1:50 mol ratio of WSN33_PB2_g2 or NTCR guides via Messenger Max. A two-ANOVA with Sidak’s multiple comparisons was performed, where * p < 0.02, ** p < 0.0024, *** p < 0.00091, and **** P < 0.0001. **c)** A549 cells were infected with IAV A/WSN/33 at MOI 0.01. 24hpi, cells were transfected with 0.5 μg Cas13a – V5 or Cas13a-V5-NLS, dCas13a-V5, dCas13a-V5-NLS mRNA or an equal molar amount of GFP mRNA (0.2 μg) with a 1:50 mol ratio of WSN33_PB2_m4 or NTCR guide via Messenger Max. A two-ANOVA with Sidak’s multiple comparisons was performed, where * p < 0.021, ** p < 0.005, and *** p < 0.00051. In all experimental conditions, cells were lysed and RNA was extracted 24 hpt (48 hpi). RT-qPCR was performed for WSN/33_PB2 by absolute copy number quantification. Means and standard deviations are shown in grey, with n = 6 biological replicates. RT-qPCR technical triplicates were performed to determine biological replicate values. WSN/33_PB2 gene copy numbers were normalized by NTCR values.

### Screening of crRNAs targeting PB2 post-infection and in combination with a PB1 guide

The mRNA targeted guides for PB1 and PB2 were then combined with both cytoplasmic and NLS targeted Cas13a and the combination was evaluated both as a function of MOI and over-time (**Fig. 6**). The use of the combination NLS and cytoplasmic Cas13a is justified by the variability of the distribution of viral RNPs from cell to cell **(Supplementary Fig 3)**. First, when the combination was evaluated, in our standard screening assay, we clearly saw the benefits of the combination, yielding ~10% lower RNA values than with WSN33_PB1_m5 alone. When this combination was examined across MOI, it was clear that at MOI=0.001 and 0.01, > than 80% knockdown was achievable, where at MOI=0.1, the effect was only ~25% **(Fig 6. a-b**). The effect though, may be increased if the enzymatic reaction is given more time, and if the dose of mRNA and guide is increased. When the combination was evaluated over time, at MOI=0.01, the results demonstrated continued RNA reduction over a span of 72 hours (**Fig 6. c-d**), with the peak difference from IAV and the NTCR at 48 hours. It should be noted that we did see increased effects of the NTCR over time, likely due to low levels of NTCR overlap with native genes inducing activation. However, if the effect of the treatment or “drug”, Cas13a mRNA+targeted guide, is compared with the IAV only case, the approach was able to sustain 90% knockdown over the 3-day period.

**Fig 6.**
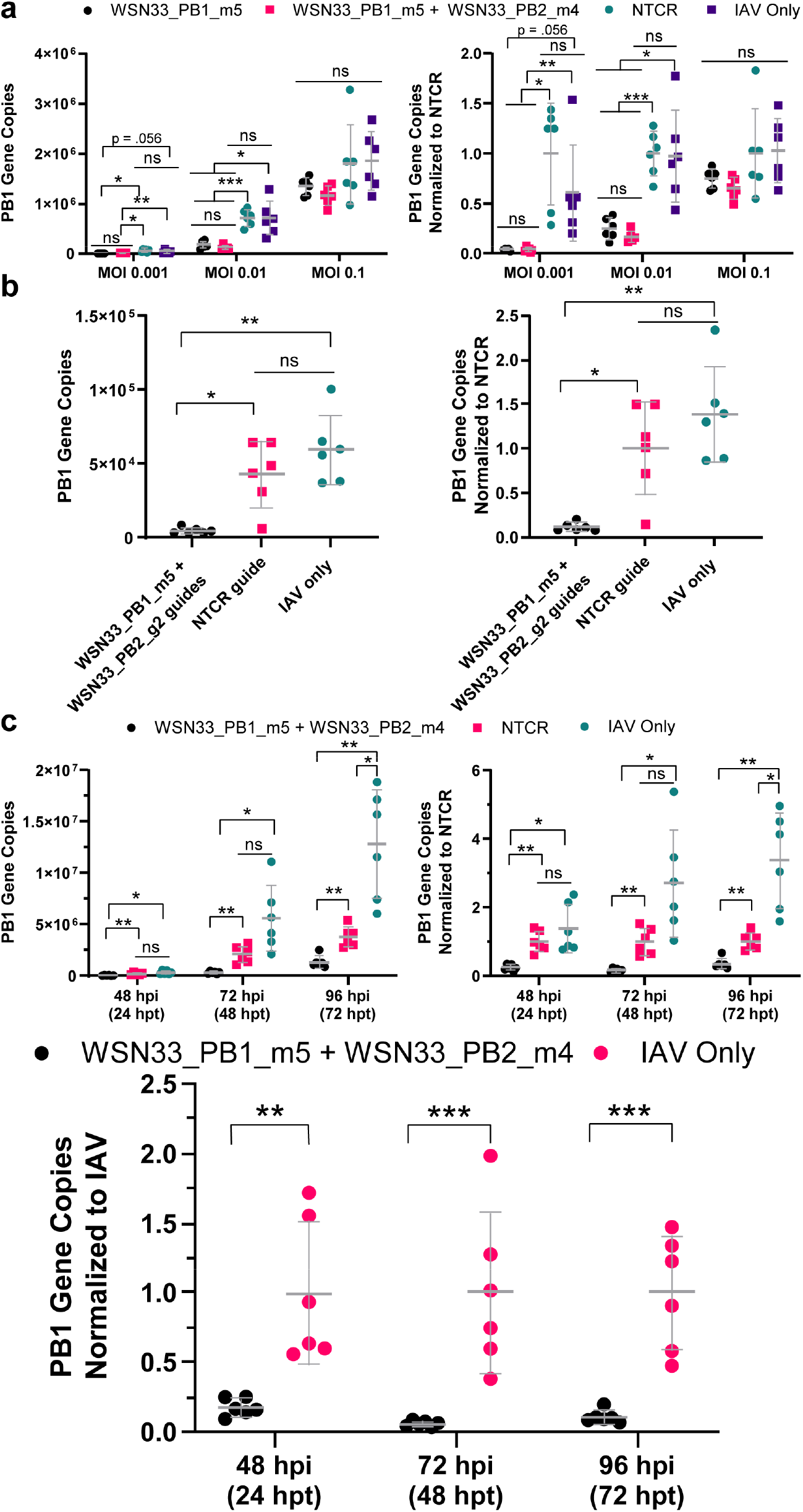
Cas13a-mediated IAV PB1 RNA knock down as a function of MOI and time-course via combinations of targeted guides. **a)** A549 cells were infected with IAV A/WSN/33 at MOI 0.001, MOI 0.01, or MOI 0.1. 24 hpi, cells were transfected with 0.25 μg of Cas13a-V5, 0.25 μg Cas13a-V5-NLS, 1:50 mol ratio of WSN33_PB1_m5 and WSN33_PB1_m5/WSN33_PB2_m4, or NTCR guides via Messenger Max. 24 hpt (48 hpi), cells were lysed and RNA was extracted. Kruskal-Wallis or Brown-Forsythe and Welch ANOVAs with Dunnett T3 multiple comparison tests were performed, where * p < 0.04, ** p < 0.01, and *** p < 0.009. **b)** A549 cells were infected with IAV A/WSN/33 at MOI 0.01. 24 hpi, cells were transfected with 0.25 μg of Cas13a-V5 and 0.25 μg Cas13a-V5-NLS and 1:50 mol ratio of WSN33_PB1_m5 and WSN33_PB2_g2, or NTCR guides via Messenger Max. 24 hpt (48 hpi), cells were lysed and RNA was extracted. Brown-Forsythe and Welch ANOVAs with Dunnett T3 multiple comparison tests were performed, where * p < 0.025, and ** p < 0.006. **c)** A549 cells were infected with IAV A/WSN/33 at MOI 0.01. 24 hpi, cells were transfected with 0.25 μg of Cas13a-V5 and 0.25 μg Cas13a-V5-NLS with a 1:50 mol ratio of WSN33_PB1_m5/WSN33_PB2_m4, or NTCR guides via Messenger Max. 24 hpt (48 hpi), 48 hpt (72 hpi), or 72 hpt (96 hpi), cells were lysed and RNA was extracted. Brown-Forsythe and Welch ANOVAs with Dunnett T3 multiple comparison tests were performed (top), where * p < 0.03, ** p < 0.01, and *** p < 0.009. A 2-way ANOVA with a Sidak’s multiple comparisons test (bottom) was performed where ** p < 0.002, and **** p < 0.0006. RT-qPCR was performed for WSN/33_PB1 by absolute copy number quantification. Means and standard deviations are shown in grey, with n = 6 biological replicates. RT-qPCR technical triplicates were performed to determine biological replicate values. WSN/33_PB1 gene copy numbers were normalized by NTCR values.

### Inhibition of SARS-CoV-2 in a Vero E6 model of infection

Given the current importance of SARS-CoV-2, a set of crRNAs that target highly conserved regions in the replicase and nucleocapsid regions of the genome were designed (**Fig. 7 and Supplementary Table 2, 3 and 4**). The nucleocapsid sites would also allow for the targeting of any of the subgenomic RNAs produced by the virus. In this experiment, 9 different guides were screened, by first transfecting Vero E6 cells overnight with the cytoplasmic version of Cas13a and guides, and then infecting with MOI=~0.1 of SARS-CoV-2. Note, only the cytoplasmic version was used, because the virus replicates in this cell compartment ^26^. Prophylactic delivery was used because this virus exhibits very rapid kinetics in Vero E6 cells, as well as the fact that they are interferon deficient. In less permissive, less cytopathic susceptible cell types, we anticipate the post-infection approach to yield similar results to the influenza experiments. At 1-hour PI, an Avicel overlay was added to the wells and CPE was evaluated at 60 hrs PI. From this screen, guides N3.2 (nucleocapsid), N3.1 (nucleocapsid), R5.1 (replicase), and N11.2 (nucleocapsid) were found to have an impact on cytopathic effect (CPE), with guide 3.2 (**Fig. 7**) exhibiting the lowest CPE. This was repeated in 6 well plates using both individual and combinations of guides including all the controls, and the data quantified by image analysis, demonstrating for the combination of N3.2 and N11.2 over 72% reduction in cell death with over 80% of cells remaining in the plate. Overall, this bodes well for use as a therapeutic for SARS-CoV-2.

**Fig 7.**
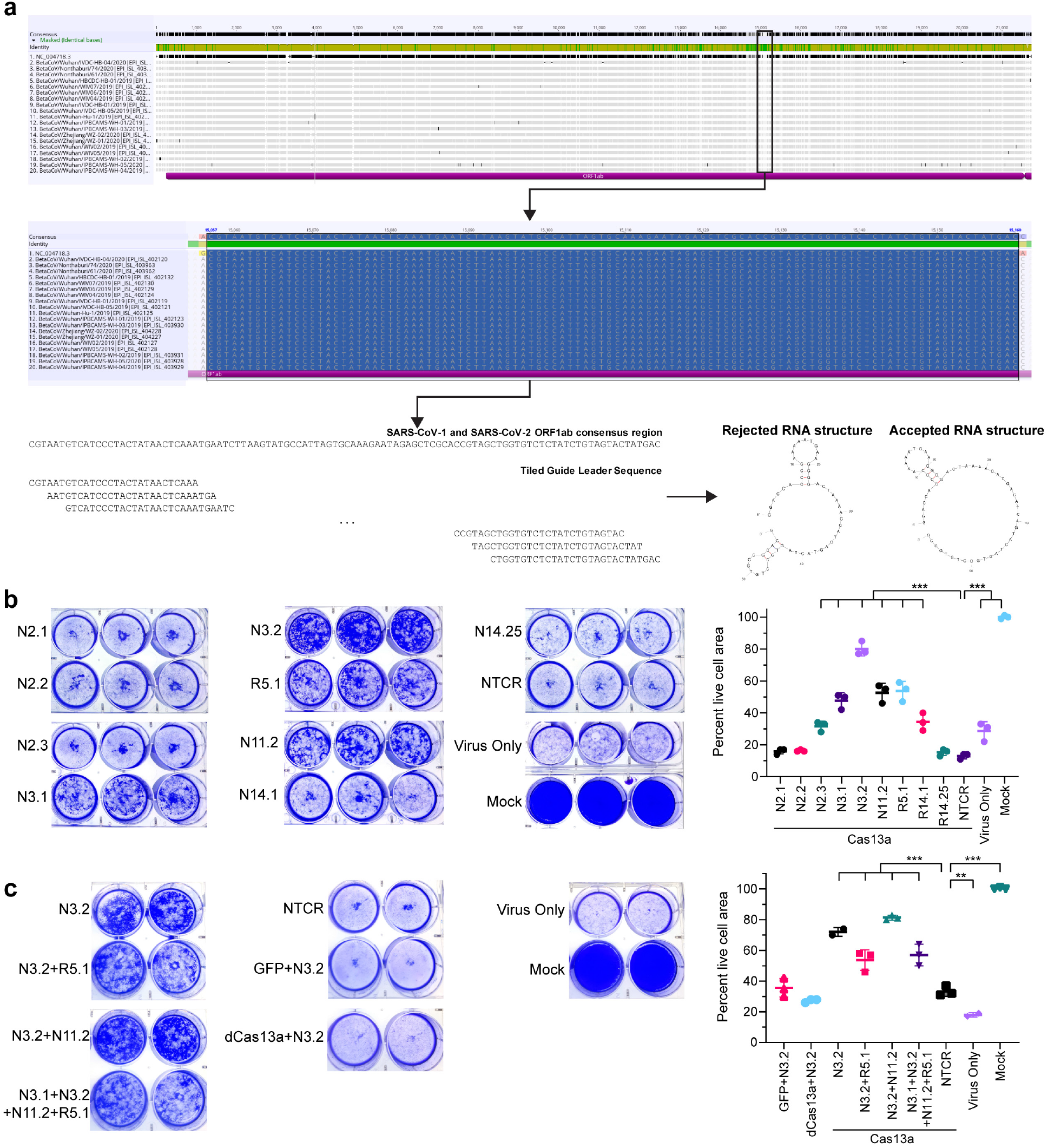
SARS-CoV-2 SARS-CoV-2 targeted guide selection and *in vitro* testing. **a)** Guide selection process for SARS-CoV-2. 19 sequences of SARS-CoV-2 from the Wuhan region were aligned with the Toronto 2 SARS-CoV isolate sequence, and regions in ORF1ab and N with complete coverage were isolated. Guide target sequences were then tiled across the conserved region and checked for correct folding. **b)** Cas13a-lbu mRNA along with each of 8 guides against N, 1 guide against ORF1ab (R5.1), or NTCR were transfected in Vero-E6 cells overnight prior to infection with SARS-CoV-2 (MOI 0.1). At 60 hpi, crystal violet stain was used to assess CPE. Percent live cell area was then plotted. Error bars indicate SD. *** indicates p≤0.0001 (one-way ANOVA with multiple comparisons to NTCR). **c)** Cas13a-lbu mRNA along with either guide N3.2 alone or in the indicated combinations was transfected in Vero-E6 cells overnight. GFP and dCas13a-lbu mRNA along with guide N3.2 was used as controls to demonstrate catalytic guide activity. Cells were infected with SARS-CoV-2 (MOI 0.1), and at 60 hpi crystal violet stain was used to assess CPE. Percent live cell area was then plotted. Error bars indicate SD. ** indicates p=0.0082; *** indicates p≤0.005 (one-way ANOVA with multiple comparisons to NTCR).

### Reduction of IAV *in vivo* post-infection

In order to test this approach *in vivo*, we first had to develop both an apparatus and formulation for the RNA for nebulizer-based delivery. We designed a straightforward nose-cone nebulizer apparatus (**Fig. 8a**), which allows for 3 mice to be dosed simultaneously from a vibrating mesh nebulizer. PBAE was chosen for our formulation based on prior publication and lung delivery^25^. First, the final concentration of mRNA in the formulation was optimized using a GPI-anchored nanoluciferase (aNLuc) encoding mRNA in the lung (**Fig. 8b**), yielding optimal expression in the lungs at 0.5 μg/ml, as reported previously. Next, we determined whether influenza infection affects delivery. We infected mice with 3 LD_50_ of Influenza A/WSN/33 and then delivered aNLuc mRNA at 12 and 24 hrs post-infection as well as without infection (**Fig. 8c-d**). No significant differences in expression were observed, demonstrating efficient mRNA delivery and protein expression via nebulizer in the face of influenza infection.

**Fig 8.**
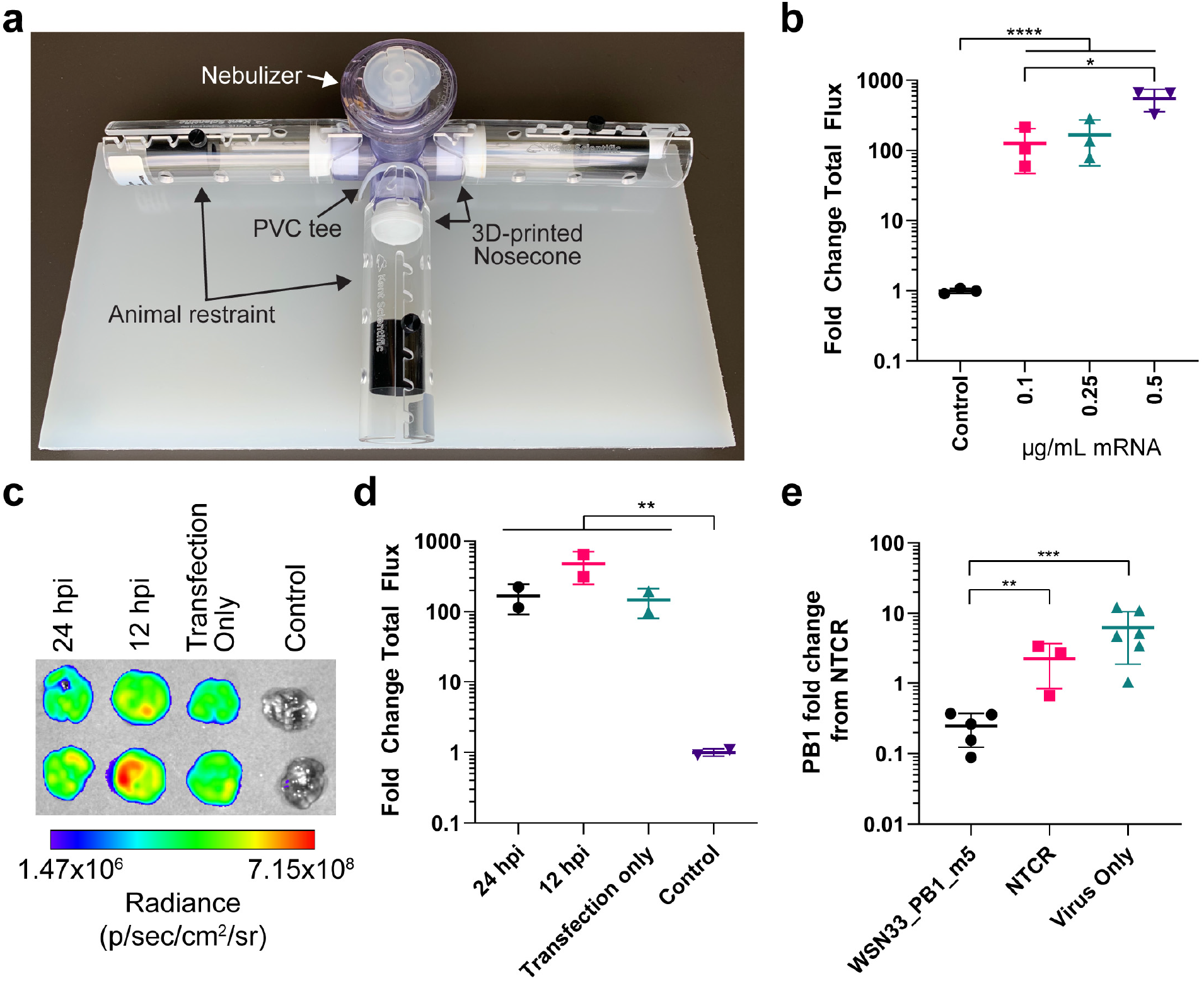
Nebulization setup and anti-influenza activity of inhalable Cas13a mRNA formulation. **a)** Custom apparatus for mouse inhalation studies. A clear PVC tee is attached with a 3D-printed nose-cone made of flexible TPU to small rodent restraints. The Aerogen nebulizer is placed into the top-facing tee slot and doses are added dropwise to the nebulizer mesh. **b)** Mice were allowed to inhale 100 μg of nebulized aNLuc mRNA formulated with PBAE at the indicated final mRNA concentration. Lungs were harvested after 1 day and analyzed for luminescence. Quantification of luminescence represented as the fold change in the total flux relative to the control lungs. Bars represent mean ± SD. **** represents p<0.0001; * indicates p<0.05 by one-way ANOVA with multiple comparisons on log-transformed data. **c)** Mice were infected with 3 LD_50_ Influenza A/WSN/33 either 12 or 24 hours, or not infected, prior to being dosed with 100 μg of nebulized aNLuc mRNA. At 1 day, lungs were harvested and analyzed for luminescence. Image is presented in log-format with the indicated radiance scale. **d)** Quantification of luminescence in part **c)** represented as the fold change of total flux. Bars represent mean ± SD. * * indicates p<0.05 by one-way ANOVA with multiple comparisons on log-transformed data. **e)** Lung viral loads from infected mice dosed with Cas13a mRNA with either targeted or NTCR guides at 6 h post infection. One group of infected mice was not treated. Data represents the mean fold change ± SD from the NTCR (left). ** represents p=0.0097; *** represents p=0.0001 by one-way ANOVA on log-transformed data.

In order to test this treatment approach, mice were infected with 3 LD_50_ of Influenza A/WSN/33 (**Fig. 8e)** via intranasal administration. Six-eight hours post-infection, one group was given, by nose cone nebulizer, 100 μg of mRNA encoding Cas13a (with and without NLS) and guide (WSN/33_PB1_m5) formulated with the PBAE-based polymer. Control groups included mice given 100 μg of Cas13a mRNA with an NTCR guide as well as an infection only group. At 3 days post-infection the animals were euthanized, and the viral RNA in their lungs was quantified by qPCR. Analysis revealed a 89.1% (p=0.0097) reduction of viral RNA from NTCR and a 96.2% (p=0.0001) from IAV only, demonstrating robust knockdown of viral RNA *in vivo*.

## DISCUSSION

RNA viruses pose challenges for drug and vaccine development and thus are a global health concern. Unconventional molecular tools like RNA-activated RNases, such as Cas13a, could be a potential new paradigm for therapeutics against these pathogens. However, for the RNA targeting RNases to be safe and effective for therapeutic use, rapid, transient expression is preferred ^27^. To achieve this, we opted for synthetic *in vitro* transcribed mRNA to express Cas13a. mRNA has the advantage of rapid translation of the desired protein and clearance, while it avoids safety concerns such as genome integration and unmitigated innate immune responses^20,21,28–30^.

Using this all RNA-based approach, we found crRNA guides that targeted both the genome and the mRNA of influenza effectively, even when given 24 hours post-infection. *To date, this is the first demonstration that this approach can be used post-infection.* There was, though, a clear bias towards mRNA targeted guides, which is consistent with prior work using siRNA against influenza^31^. In addition, even though we were not able to find guides that were truly pan-influenza, they clearly overlap with over 52,000 strains, making significant strides towards this goal. Prior studies had not targeted polymerase genes. It is our contention that they are excellent targets due to their clear role in generating viral RNA and the fact they are so well conserved. *We also demonstrated for the first time, how this approach scales with MOI and is functional over a 3-day period even when using transient transfection of mRNA.* The time course data demonstrated the ability of Cas13a to degrade RNA at *a rate equal to or more rapidly* than the rate of RNA generation from viral replication. This has not been addressed to date. The data clearly shows that over the 3-day period post-delivery, that Cas13a was keeping RNA levels consistently ~1 log lower than without the treatment. In the future, we will address how dose effects these dynamics and how this may change with Cas13b or d.

We then addressed the question; can this approach be rapidly applied directly to newly emerging pathogens like SARS-CoV-2? From our results, it is clear that Cas13a can have *significant effects on CPE*, a well-established measure of antiviral activity^32^. These results clearly demonstrate the adaptability of this approach and its flexibility. Different guide sequences are sufficient to target either IAV or SARS-CoV-2. By delivering the nuclear or the cytoplasmic versions of Cas13, or both together, the replication and dynamics of different viruses can be significantly impacted. Given the promising results *in vitro* for SARS-CoV-2, this approach will be moved into animal models in the very near future. Due to the need for ABSL3, and the fact that the animal models are not yet mature, we did not pursue these experiments at this juncture.

Last, we answered a critical question regarding Cas13, can it be used *in vivo*? Here we formulated the mRNA with PBAE-based polymer previously demonstrated by the Anderson lab^25^ and delivered it post-infection using a vibrating-mesh style nebulizer that is currently used by humans, simulating a treatment strategy, and demonstrated that Cas13 can degrade viral RNA *in vivo* efficiently. This is the first demonstration of Cas13 use in an animal model of infection and it bodes well for treating other infections, such as SARS-CoV-2. Given we have guides that were effective *in vitro*, we have confidence that we can move this to an appropriate animal model such as ferrets or hamsters. This will be a very important step towards developing a Cas13 based treatment strategy for SARS-CoV-2. Future work will include these vital steps in addition to further development of mRNA carriers for the lung. Overall, this work demonstrates significant progress toward the use of this of Cas13 to treat respiratory viral infections.

## METHODS

### Design and synthesis of Cas13a constructs and anchored nanoluciferase mRNA

The Cas13a sequence from *Leptotrichia buccalis* was obtained from Addgene p2CT-His-MBP-Lbu_C2C2_WT (Plasmid #83482). We cloned the wild type Cas13a with a 3’ V5 tag (Cas13a-V5) and appended a 3’ UTR from mouse alpha-globin (Genbank accession # NM_001083955) in a pMA7 vector (GeneArt, Thermo Scientific, USA). Additionally, we synthesized a construct with the Cas13a sequence from the *Leptotrichia buccalis* along with a nuclear localization sequence (at 3’ and 5’) and a 3’ V5 tag to create a Cas13a-V5-NLS version using GeneBlocks (Integrated DNA technologies). For both constructs, we synthesized catalytically-inactive versions (dCas13a – V5 and dCas13a-V5-NLS). The sequences were obtained by Addgene plasmid#100817. The sequences of the GPI anchor and nanoluciferase are described in Lindsay et al^33^.

### *In vitro* transcription of Cas13a

Plasmids were linearized with Not-I HF (New England Biolabs) overnight at 37°C. Linearized templates were purified by sodium acetate (Thermo Fisher Scientific) precipitation and rehydrated with nuclease free water. IVT was performed overnight at 37°C using the HiScribe T7 kit (NEB) following the manufacturer’s instructions (N1-methyl-pseudouridine modified). The resulting RNA was treated with DNase I (Aldevron) for 30 min to remove the template, and it was purified using lithium chloride precipitation (Thermo Fisher Scientific). The RNA was heat denatured at 65°C for 10 minutes before capping with a Cap-1 structure using guanylyl transferase and 2’-O-methyltransferase (Aldevron). mRNA was then purified by lithium chloride precipitation, treated with alkaline phosphatase (NEB) and purified again. mRNA concentration was measured using a Nanodrop. mRNA stock concentrations were 1-3 mg/ml. Purified mRNA products were analyzed by gel electrophoresis to ensure purity. crRNA guides were purchased from Integrated DNA Technologies (IDT) or Genscript. The sequences are detailed in **Supplementary Table 2**.

### RNA cleavage activity of *in vitro* translated of Cas13a

In order to demonstrate that mRNA-expressed Cas13a efficiently encodes for a fully functional enzyme able to cleave a target RNA in a “in tube” assay, Cas13a mRNA was translated *in vitro* using the Rabbit reticulocyte lysate system according to the manufacturer’s instructions (Promega, WI, USA). RNaseAlert®-1 Substrate was mixed with crRNA or NTCR (500 nM) and trRNA (500 nM) in RNA processing buffer, consisting of A549 cells RNA (100 ng), 20 mM HEPES (pH 6.8), 50 mM KCl, 5 mM MgCl_2_, BSA (10 μg/ml), yeast tRNA (10 μg/ml), 0.01% Igepal CA-630 and 5% glycerol (Supplementary Table 1)^15^. This mixture was added cold to the translated Cas13a lysate (5 μl) in the 96-well plate wells and mixed well. All the reagent preparations and additions were performed on ice. The fluorescence measurements (excitation 485±20 nm/emission 528 ± 20 nm) were recorded at room temperature for 90 min at 10 min interval.

### Cell lines and viruses

All cell lines and viruses were purchased from American Type culture collection (ATCC. Manassas, VA). Human lung epithelial cells A549 (CCL185), MDCK, Vero E6 were grown in media recommended by ATCC. Guide screening experiments for IAV were performed in A549 cells. Guide screening experiments for SARS-CoV-2 were performed in Vero E6 cells. Influenza virus stocks (H1N1 Influenza virus A/WSN/33) were prepared in MDCK cells. Briefly, MDCK cells were grown till 100% confluence in 175 mm^2^. The next day, cells were washed twice with PBS and 1:1000 dilution of virus was added in 5 ml EMEM. Cells were then incubated with the virus for 1h at room temperature on a rocker. Then, 25 ml media was added to the cells. Cells were monitored for 72 h or until a severe cytopathic effect was observed. Virus was collected by centrifuging the cells at 1000xg for 10 min. Virus titers were determined by standard plaque assay.

SARS-CoV-2 (USA-WA1/2020) was obtained from BEI Resources (Manassas, VA). Viral stocks were generated by infecting Vero E6 cells (ATCC, C1008) at ~95% confluency in 150cm^2^ flasks with SARS-CoV-2 at an MOI of 0.1 PFU/cell. At 68 hours post-infection, supernatants were collected, pooled, and centrifuged at 400*xg* for 10 minutes. The resulting stock was aliquoted, titered, and stored at −80°C for further use. All work with live SARS-CoV-2 was performed inside a certified Class II Biosafety Cabinet in a BSL3 laboratory in compliance with all State and Federal guidelines and with the approval of the UGA Institutional Biosafety Committee (IBC).

### Optimization of mRNA transfection

A549 cells were seeded overnight at a density of 120,000-130,000 per well in a 24 well plate. The following day, cells were transfected with mRNA encoding for Cas13a and either targeted crRNA or non-targeted crRNA guides (NTCR) using Lipofectamine Messenger Max (Thermo Fisher Scientific) according to manufacturer instructions. dCas13 constructs-encoding mRNAs and GFP-encoding mRNA were also used as controls to assess the effect of Cas13a targeted binding and mRNA transfection/guides alone, respectively, on gene expression. For each well, 0.5 μg mRNA and 1.5ul Messenger Max were used. crRNA guides and NTCR were added to the transfection mix at 20x molar guide:mRNA ratio in all endogenous gene knockdown experiments (PPIB, CXCR4, KRAS). The sequences of the previously published crRNA guides^10^ are detailed in **Supplementary Tables 2 and 3**.

24h post transfection, total RNA was extracted using RNeasy Plus mini kit. cDNA was prepared using the High-Capacity cDNA reverse transcription kit (The Applied Biosystems™, Thermo Fisher Scientific). qPCR experiments were performed using the FastAdvanced Master Mix (Thermo Fisher Scientific). Cas13-mediated degradation of PPIB, CXCR4 and KRAS mRNA was assayed by qPCR with the ΔΔCT method (n=3) using GAPDH as control. Experiments were performed using a QuantStudio7 Flex thermal cycler (Applied Biosystems). All primer/probe assays for endogenous gene experiments were purchased from Thermo Fisher Scientific **(Supplementary Table 5).**

### *In vitro* anti-viral assay with IAV

A549 cells were seeded overnight at a density of 120,000-130,000 per well in a 24 well plate. The following day, cells were washed with 1XPBS, and infected by adding 100ul of Influenza virus diluted in serum-free media with TPCK for 1h. The cells were then washed with 1xPBS and complete media with TPCK was added for 24h. Cells were then transfected with mRNA encoding for Cas13a and either targeted crRNA or non-targeted crRNA guides (NTCR) using Lipofectamine Messenger Max (Thermo Fisher Scientific) according to manufacturer instructions. dCas13 constructs-encoding mRNAs and GFP-encoding mRNA were also used as controls to assess the effect of Cas13a targeted binding and mRNA transfection/guides alone, respectively, on gene expression. For each well, 0.5 μg mRNA and 1.5ul Messenger Max were used. crRNA guides and NTCR were added to the transfection mix at 50x molar guide:mRNA ratio in all IVA-based experiments. The sequences of the crRNA guides are detailed in **Supplementary Tables 2 and 3**.

24h post transfection, total RNA was extracted using RNeasy Plus mini kit. cDNA was prepared using the High-Capacity cDNA reverse transcription kit (The Applied Biosystems™, Thermo Fisher Scientific). qPCR experiments were performed using the FastAdvanced Master Mix (Thermo Fisher Scientific). The anti-viral activity of Cas13a system was measured by qPCR (n=6) with the absolute quantification of the viral PB1 or PB2 gene copy number. Experiments were performed using a QuantStudio7 Flex thermal cycler (Applied Biosystems).

In the “combo experiments”, for each well, 0.25 μg of each mRNA Cas13a – V5 and Cas13a-V5-NLS and 1.5ul Messenger Max were used. Each crRNA guides was added to the transfection mix at 25x molar guide:mRNA ratio. The sequences of the primer/probe assays for PB1 and PB2 for H1N1 Influenza virus A/WSN/33 are listed in **Supplementary Table 5**.

### *In vitro* anti-viral assay with SARS-CoV-2

Vero cells were seeded overnight at ~80% confluency in a 6 well plate. The following day, cells were transfected with mRNA encoding for Cas13a-V5 and either targeted crRNA or non-targeted crRNA guides (NTCR) using Lipofectamine Messenger Max (Thermo Fisher Scientific) according to manufacturer instructions. For each well, 5 μg mRNA and 7.5ul Messenger Max were used. crRNA guides and NTCR were added to the transfection mix at 50x molar guide:mRNA ratio (**Supplementary Tables 2 and 3**). In the “combo experiments”, for each well, 5 μg of Cas13a – V5 and 7.5ul Messenger Max were used. Each crRNA guide was added to the transfection mix to maintain the 50x molar guide:mRNA ratio. After an overnight incubation at 37 °C, 5% CO2, plates were transferred to the BSL3 for infection. The medium was removed, and cells were infected with SARS-CoV-2 for 45-60 minutes at an MOI of ~0.1 PFU/cell. After infection, cells were overlayed with 1x DMEM 1%FBS containing 1.2% Avicel RC-581 and incubated for 72 hours. The overlay was removed, cells rinsed with 1x PBS, and fixed/stained with a crystal violet solution containing 2% methanol, 4% formaldehyde for 10 minutes. Images for presentation were white balanced and intensities for analysis were not modified for calculations. All images were brought into photoshop, converted to grayscale, and color inverted. An ROI of the same size was then used for all images and the sum intensity was calculated by Volocity. Intensities were normalized to a blank region of each image in order to account for image to image variation. Signal was then normalized to the Mock condition to give a percent live cell area.

### Animal studies

Six- to 8-week-old female BALB/c mice (Jackson Laboratories) were maintained under pathogen-free conditions in individually ventilated and watered cages kept at negative pressure. Food was provided to mice *ad libitum*. Animals were acclimatized for at least 6 days before the beginning of experiments. Animals were randomly distributed among experimental groups. Researchers were blinded to animal group allocation during data acquisition. Animals were sacrificed by CO_2_ asphyxiation. Infected animals were handled and kept under BSL-2 conditions until euthanized. All animals were cared for according to the Georgia Institute of Technology Physiological Research Laboratory policies and under ethical guidance from the university’s Institutional Animal Care and Use Committee (IACUC) following National institutes of Health (NIH) guidelines.

### Nebulizer-based mRNA deliveries

Mice were loaded into a custom-built nose-only exposure system constructed of a clear PVC tee and animal restraints (CODA Small Mouse Holder, Kent Scientific). These were connected using a custom 3D-printed nosecone (3D Printing Tech) made of a flexible TPU material. The nebulizer (Aeroneb, Kent Scientific) was then placed on the upward facing port of the tee. Doses were added dropwise to the nebulizer at a rate of 25 μL/mouse/droplet. After each individual droplet was nebulized, the clear tee was inspected until the vaporized dose had cleared (approximately 15-45 seconds per drop). Droplets were added until the desired dose per animal was achieved. After the vapor had cleared following the last droplet, the mice were removed from the restraints.

mRNA was formulated for nebulizer-based delivery using a hyperbranched PBAE as previously described^25^ with minor modifications. Diacrylate and amine monomers were purchased from Sigma-Aldrich. To synthesize hyperbranched hDD90-118, acrylate: backbone amine: trifunctional amine monomers were reacted modifying the ratio at 1: 0.59: 0.40. Monomers were stirred in anhydrous dimethylformamide at a concentration of 150 mg/mL at 40 °C for 4 h then 90 °C for 48 h. The mixtures were allowed to cool to 30 °C and end cap amine was added at 1.0 molar equivalent relative to the acrylate and stirred for a further 24 h. The polymers were purified by dropwise precipitation into cold anhydrous diethyl ether spiked with glacial acetic acid, vortexed and centrifuged at 1250 xg for 2 min. The supernatant was discarded, and the polymer washed twice more in fresh diethyl ether and dried under vacuum for 48 h. This step was repeated until the supernatant in the precipitation process became transparent. Polymers were stored at −20 °C.

Before delivery to mice, 100 mM of sodium acetate pH 5.0 was used to both solubilize the hyperbranched PBAE and dilute mRNA prior to mixing. The final concentration of the mRNA was 0.5 mg/mL, and the PBAE was used at a 50x molar ratio to the mRNA. The tubes were incubated at RT for 10 minutes, and the particles were loaded into the nebulizer as described above.

Following euthanasia, whole lungs were collected and rinsed with PBS. Lungs were then placed into a solution of Nano-Glo substrate solution (Promega) diluted 50-fold in PBS. Lungs were incubated for 5 minutes and then placed onto black paper and imaged with an IVIS Spectrum CT (Perkin Elmer). Lung luminescence was then quantified using Living Image software (Perkin Elmer).

### Viral load quantification from lung tissues

Mice were infected intranasally with 3 LD_50_ influenza virus using 20 μL per nostril following brief anaesthesia by isoflurane. Virus was diluted in DMEM without modifications to the desired PFU dose. After euthanasia, lungs were harvested into PBS prior to downstream assays.

Lungs were dissociated as described previously^29^. Briefly, lungs were weighed before dissociation using a GentleMACS in a C Tube using the lung 2 setting (Miltenyi). After centrifugation at RT for 5 minutes at 500 xg, pellets were lysed in RLT plus buffer (Qiagen) and further homogenized using NAVY tubes in a Bullet Blender (Next Advance). Lysates were centrifuged for 15 min at 4 °C and maximum speed to clarify the supernatant. Total RNA was then extracted following a RNeasy Plus Mini kit (Qiagen) per manufacturer’s instructions. cDNA synthesis and qPCR were then performed as detailed above with the absolute quantification of the viral PB1 gene copy number.

### Statistical Analyses

All experiments are represented as a mean of three or six biological replicates as indicated. Data was analyzed using GraphPad Prism 7.04. Statistical analyses were performed between groups using either ordinary one-way or two-way analysis of variance (ANOVA) as specified in individual figure captions.

## AVAILABILITY OF DATA

All data generated or analysed during this study are included in this published article and its supplementary information files.

## AUTHOR CONTRIBUTION

ELB SB CZ and PJS Developed screening assays, ELB Performed PB1 guide screens, guide combinations, MOI and time course studies *in vitro*. DV Developed nose-cone nebulizer setup and optimized polymer-based delivery, performed *in vivo* experiments and data analysis, designed SARS-CoV-2 guides and transfections. SB and PT designed Cas13 mRNA and guides and PT performed PB2 screens. CZ performed endogenous gene knockdown experiments. LR synthesized the polymer and assisted with *in vivo* studies. NB, RH, and MGF assisted with polymer production. JB assisted with delivery experiment. HP synthesized mRNA and assisted with *in vivo* studies. JH, FM, and ERL performed SARS-CoV-2 CPE assays. PJS designed experiments and wrote the manuscript.

## ACKNOWLEDGEMENT

The authors would like to acknowledge Ian A. York from the Influenza Division, National Center for Immunization and Respiratory Diseases, Centers for Disease Control and Prevention, Atlanta, GA USA for providing the influenza and SARS-CoV-2 sequences used in this manuscript. The following reagent was deposited by the Centers for Disease Control and Prevention and obtained through BEI Resources, NIAID, NIH: SARS-Related Coronavirus 2, Isolate USA-WA1/2020, NR-52281.

---

**Fig S1 Cas13a-mediated IAV PB1 RNA knock down post-infection. (additional graphs). a)** A549 cells were infected with IAV A/WSN/33 at MOI 0.01. 24 hpi, cells were transfected with 0.5 μg of Cas13a-V5 or Cas13a-V5-NLS and 1:50 mol ratio of WSN33_PB1 or NTCR guides via Messenger Max. Two-ANOVAs with Sidak’s multiple comparisons were performed, where ** p < 0.009, *** p = 0.0001, and **** p < 0.0001. **b)** A549 cells were infected with IAV A/WSN/33 at MOI 0.01. 24 hpi, cells were transfected with 0.5 μg Cas13a – V5 or Cas13a-V5-NLS, dCas13a-V5, dCas13a-V5-NLS mRNA or an equal molar amount of GFP mRNA (0.2 μg) with a 1:50 mol ratio of WSN33_PB1_g2 or NTCR guides via Messenger Max. A two-ANOVAs with Sidak’s multiple comparisons was performed, where no significant differences were found. **c)** A549 cells were infected with IAV A/WSN/33 at MOI 0.01. 24 hpi, cells were transfected with 0.5 μg Cas13a – V5 or Cas13a-V5-NLS, dCas13a-V5, dCas13a-V5-NLS mRNA or an equal molar amount of GFP mRNA (0.2 μg) with a 1:50 mol ratio of WSN33_PB1_m5 or NTCR guides via Messenger Max. A two-ANOVA with Sidak’s multiple comparisons was performed, where *** p < 0.00065, and **** p < 0.0001. In all experimental conditions, cells were lysed and RNA was extracted 24 hpt (48 hpi). RT-qPCR was performed for WSN/33_PB1 by absolute copy number quantification. Means and standard deviations are shown in grey, with n = 6 biological replicates. RT-qPCR technical triplicates were performed to determine biological replicate values.

**Fig S2- IAV PB2 RNA can be knocked down post-infection with Cas13a with broadly targeted guide RNAs (additional graphs). a)** A549 cells were infected with IAV A/WSN/33 at MOI 0.01. 24 hpi, 24 hpi, cells were transfected with 0.5 μg of Cas13a-V5 or Cas13a-V5-NLS and 1:50 mol ratio of WSN33_PB2 or NTCR guides via Messenger Max. Two-ANOVAs with Tukey’s multiple comparisons were performed, where ** p = 0.0013, and **** p < 0.0001. **b)** A549 cells were infected with IAV A/WSN/33 at MOI 0.01. 24 hpi, cells were transfected with 0.5 μg Cas13a – V5 or Cas13a-V5-NLS, dCas13a-V5, dCas13a-V5-NLS mRNA or an equal molar amount of GFP mRNA (0.2 μg) with a 1:50 mol ratio of WSN33_PB2_g2 or NTCR guides via Messenger Max. A two-ANOVA with Tukey’s multiple comparisons was performed, where * p = 0.014, ** p = 0.0059, *** p < 0.00071, and **** P < 0.0001. **c)** A549 cells were infected with IAV A/WSN/33 at MOI 0.01. 24 hpi, cells were transfected with 0.5 μg Cas13a – V5 or Cas13a-V5-NLS, dCas13a-V5, dCas13a-V5-NLS mRNA or an equal molar amount of GFP mRNA (0.2 μg) with a 1:50 mol ratio of WSN33_PB2_m4 or NTCR guides via Messenger Max. A two-ANOVA with Tukey’s multiple comparisons was performed, where ** p = 0.0028, and *** p < 0.00051. In all experimental conditions, 24 hpt (48 hpi), cells were lysed and RNA was extracted. RT-qPCR was performed for WSN/33_PB2, by absolute copy number quantification. Means and standard deviations are shown in grey, with n = 6 biological replicates. RT-qPCR technical triplicates were performed to determine biological replicate values.

**Fig S3 IAV NP localization is heterogenous during infection.** A549 cells were infected with IAV A/WSN/33 at MOI 0.001, MOI 0.01, or MOI 0.1. Cells were fixed 24 hpi and immunostained for IAV NP protein (green) and DAPI (blue). Representative single plane images are shown, with a 10 μm scale bar.

