## Supplementary Figures and Tables for "Treating Influenza and SARS-CoV-2 via mRNA-encoded Cas13a"

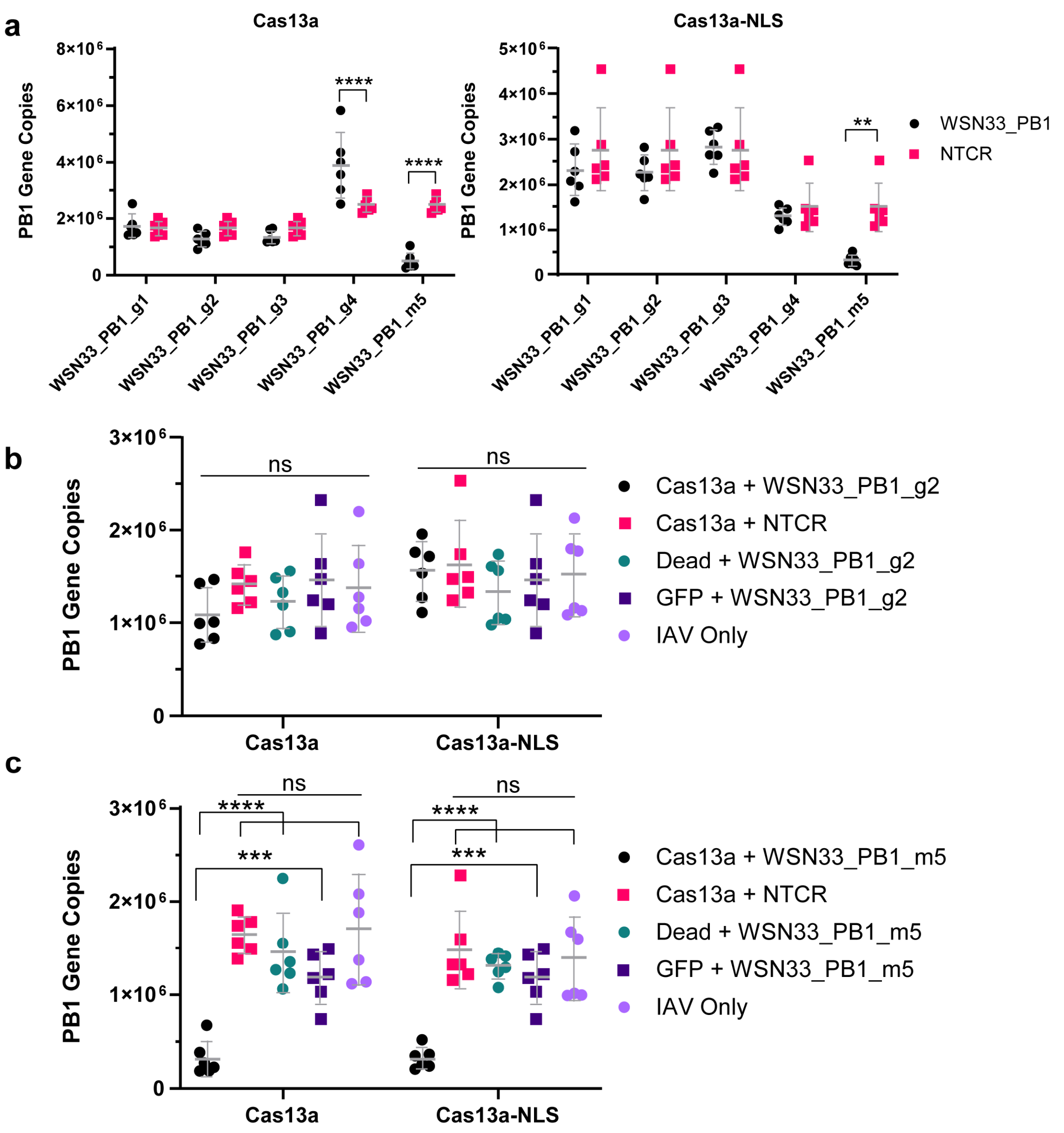
a
Cas13a
Cas13a-NLS
PB1 Gene Copies
8×10^6
6×10^6
4×10^6
2×10^6
0
5×10^6
4×10^6
3×10^6
2×10^6
1×10^6
****
**
WSN33_PB1_g1
WSN33_PB1_g2
WSN33_PB1_g3
WSN33_PB1_g4
WSN33_PB1_m5
WSN33_PB1
NTCR
b
3×10^6
2×10^6
1×10^6
ns
Cas13a + WSN33_PB1_g2
Cas13a + NTCR
Dead + WSN33_PB1_g2
GFP + WSN33_PB1_g2
IAV Only
Cas13a
Cas13a-NLS
c
***
Cas13a + WSN33_PB1_m5
Dead + WSN33_PB1_m5
GFP + WSN33_PB1_m5

Fig. S1

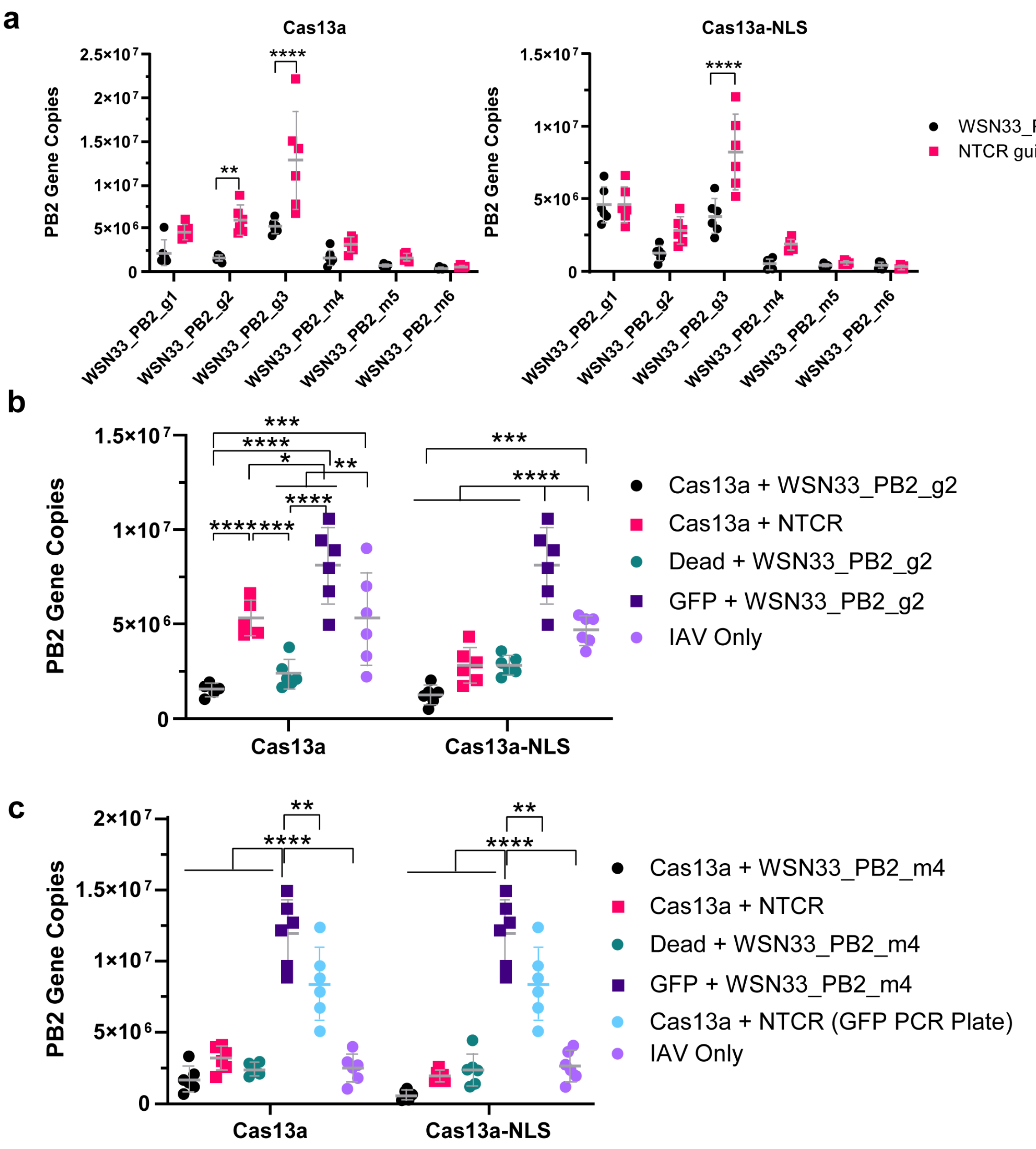
a
Cas13a
Cas13a-NLS
PB2 Gene Copies
WSN33_PB2 guide
NTCR guide
WSN33_PB2_g1
WSN33_PB2_g2
WSN33_PB2_g3
WSN33_PB2_m4
WSN33_PB2_m5
WSN33_PB2_m6
b
PB2 Gene Copies
Cas13a + WSN33_PB2_g2
Cas13a + NTCR
Dead + WSN33_PB2_g2
GFP + WSN33_PB2_g2
IAV Only
Cas13a
Cas13a-NLS
c
PB2 Gene Copies
Cas13a + WSN33_PB2_m4
Cas13a + NTCR
Dead + WSN33_PB2_m4
GFP + WSN33_PB2_m4
Cas13a + NTCR (GFP PCR Plate)
IAV Only
Cas13a
Cas13a-NLS

**Fig. S2**

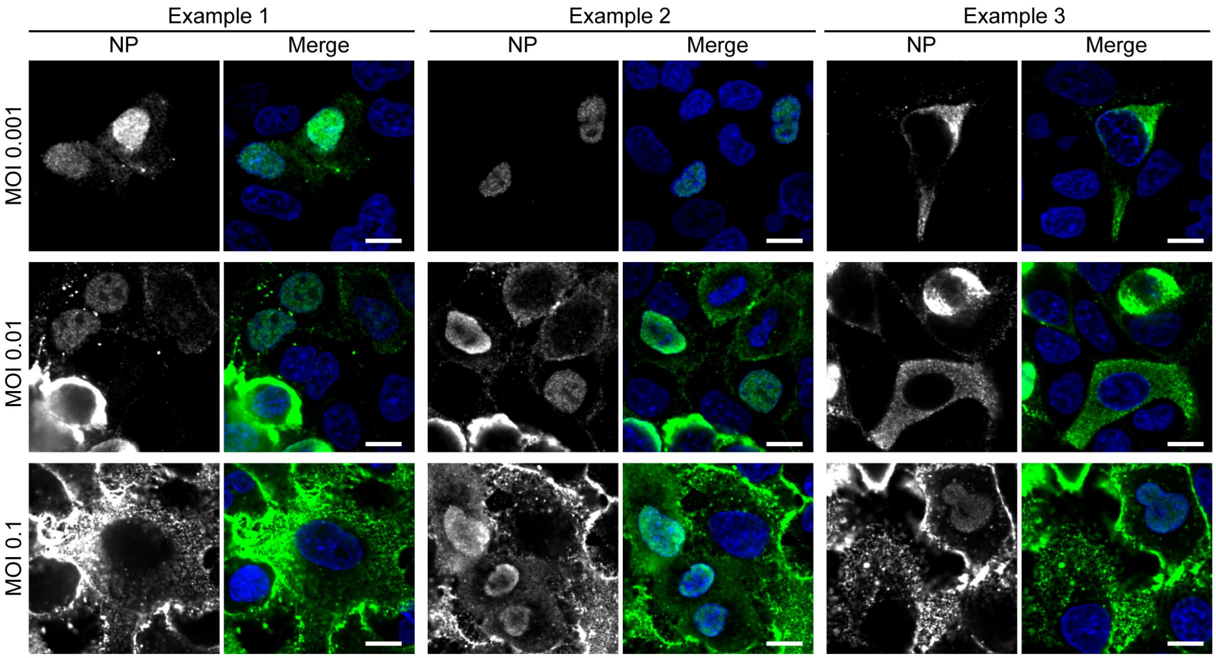
Example 1
Example 2
Example 3
NP
Merge
NP
Merge
NP
Merge
MOI 0.001
MOI 0.01
MOI 0.1

**Fig. S3**

| | |
|---|---|
| **crRNA** | 5’- GGACCACCCCAAAAAUGAAGGGGACUAAAACAAACAGAUUGUGUAUUGGAAGCAA-3’ |
| **trRNA** | 5’-GUUGCUUCCAAUACACAAUCUGUUUC-3’ |
| **NTCR** | 5’-GGUAGACCACCCCAAAAUGAAGGGGACUAAAACACAAAUCUAUCUGAAUAAACUCUUCUUC-3’ |

**Supplementary Table 1:** Sequences of crRNA, target (trRNA) and NTCR used for the *in vitro* translation assay.

| crRNA | Sequence |
|---|---|
| **Endogenous genes** | |
| PPIB* | **GGACCACCCCAAAAAUGAAGGGGACUAAAAC**UCCUUGAUUACACGAUGGAAUUUGCUGU |
| CXCR4* | **GGCCACCCCAAAAAUGAAGGGGACUAAAAC**AUGAUAAUGCAAUAGCAGGACAGGAUGA |
| KRAS* | **GGACCACCCCAAAAAUGAAGGGGACUAAAAC**AAUUUCUCGAACUAAUGUAUAGAAGGCA |
| **IAV PB1** | |
| PB1_g1 | **GGACCACCCCAAAAAUGAAGGGGACUAAAAC**AGGCCCAAAUUUAUACAACAUUAGAAAU |
| PB1_g2 | **GGACCACCCCAAAAAUGAAGGGGACUAAAAC**GCCCAAAUUUAUACAACAUUAGAAAUCU |
| PB1_g3 | **GGACCACCCCAAAAAUGAAGGGGACUAAAAC**ACGGAGGCCCAAAUUUAUACAACAUUA |
| PB1_g4 | **GGACCACCCCAAAAAUGAAGGGGACUAAAAC**AAACAGA UUGUGUAUUGGAAGCAA |
| PB1_m5 | **GGACCACCCCAAAAAUGAAGGGGACUAAAAC**UGGAGAUUUCUAAUGUUGUAUAAAUUU |
| **IAV PB2** | |
| PB2_g1 | **GGACCACCCCAAAAAUGAAGGGGACUAAAAC**GACAGCCAGACAGCGACCAAAAGAATTC |
| PB2_g2 | **GGACCACCCCAAAAAUGAAGGGGACUAAAAC** GACAGCCAGACAGCGACCAAAAGAATT |
| PB2_g3 | **GGACCACCCCAAAAAUGAAGGGGACUAAAAC**ACAGCCAGACAGCGACCAAAAGAATTC |
| PB2_m4 | **GGACCACCCCAAAAAUGAAGGGGACUAAAAC**CCGAATTCTTTTGGTCGCTGTCTGGCT |
| PB2_m5 | **GGACCACCCCAAAAAUGAAGGGGACUAAAAC**GAATTCTTTTGGTCGCTGTCTGGCTGTC |
| PB2_m6 | **GGACCACCCCAAAAAUGAAGGGGACUAAAAC**TTCTTTTGGTCGCTGTCTGGCTGTCA |
| **SARS2-CoV** | |
| N2.1 | **GGACCACCCCAAAAAUGAAGGGGACUAAAAC**GGTAGCTCTTCGGTAGTAGCCAATTTG |
| N2.2 | **GGACCACCCCAAAAAUGAAGGGGACUAAAAC**GCTCTTCGGTAGTAGCCAATTTGGTCA |
| N2.3 | **GGACCACCCCAAAAAUGAAGGGGACUAAAAC**GGTAGTAGCCAATTTGGTCATCTGGAC |
| N3.1 | **GGACCACCCCAAAAAUGAAGGGGACUAAAAC**GGTGTATTCAAGGCTCCCTCAGTTGCA |
| N3.2 | **GGACCACCCCAAAAAUGAAGGGGACUAAAAC**GTATTCAAGGCTCCCTCAGTTGCAACC |
| N3.3 | **GGACCACCCCAAAAAUGAAGGGGACUAAAAC**TCAAGGCTCCCTCAGTTGCAACCCATA |
| N7.1 | **GGACCACCCCAAAAAUGAAGGGGACUAAAAC**GCCTCAGCAGCAGATTTCTTAGTGACA |
| N10.1 | **GGACCACCCCAAAAAUGAAGGGGACUAAAAC**CGAAGGTGTGACTTCCATGCCAATGCG |
| N11.2 | **GGACCACCCCAAAAAUGAAGGGGACUAAAAC**CTGTTGGTGGGAATGTTTTGTATGCGT |
| R5.1 | **GGACCACCCCAAAAAUGAAGGGGACUAAAAC**GGTAGCTCTTCGGTAGTAGCCAATTTG |
| R14.1 | **GGACCACCCCAAAAAUGAAGGGGACUAAAAC**GCTCTTCGGTAGTAGCCAATTTGGTCA |
| R14.14 | **GGACCACCCCAAAAAUGAAGGGGACUAAAAC**GGTAGTAGCCAATTTGGTCATCTGGAC |
| R14.25 | **GGACCACCCCAAAAAUGAAGGGGACUAAAAC**GGTGTATTCAAGGCTCCCTCAGTTGCA |
| NTCR | **GGUAGACCACCCCAAAAUGAAGGGGACUAAAAC**ACAAAUCUAUCUGAAUAAACUCUUCUUC |

**Supplementary Table 2: Sequences of crRNA guides**. PPIB, KRAS multiplexing guide, CXCR4 WT guides previously published[1]. The Lbu Cas13a header sequence is indicated in bold.

| crRNA | ΔG Whole Sequence | ΔG Target sequence |
|---|---|---|
| **Endogenous genes** | | |
| PPIB* | -2.74 | 1.04 |
| CXCR4* | -4.04 | 0.89 |
| KRAS* | -1.89 | 0.89 |
| **IAV PB1** | | |
| PB1_g1 | -2.74 | 1.75 |
| PB1_g2 | -2.74 | 1.07 |
| PB1_g3 | -2.74 | 1.07 |
| PB1_g4 | -2.65 | 0.09 |
| PB1_m5 | -2.74 | 0.85 |
| **IAV PB2** | | |
| PB2_g1 | -2.74 | 0.89 |
| PB2_g2 | -2.74 | 0.89 |
| PB2_g3 | -2.74 | 0.89 |
| PB2_m4 | -2.74 | 0.6 |
| PB2_m5 | -2.74 | 0.6 |
| PB2_m6 | -2.74 | 0.6 |
| **SARS2-CoV** | | |
| N2.1 | -2.74 | 0.41 |
| N2.2 | -3.03 | 0.07 |
| N2.3 | -2.79 | -0.05 |
| N3.1 | -2.74 | 0.46 |
| N3.2 | -2.74 | 0.46 |
| N3.3 | -2.74 | 0.46 |
| N7.1 | -2.76 | 0.15 |
| N10.1 | -4.46 | -0.05 |
| N11.2 | -4.36 | 1.71 |
| R5.1 | -2.74 | 0.32 |
| R14.1 | -2.74 | 0.18 |
| R14.14 | -2.74 | 0.41 |
| R14.25 | -2.74 | 1.34 |
| NTCR | -2.74 | 1.16 |

**Supplementary Table 3**: ΔG values for all guides used in the experiments. Values are reported for the whole sequence (including header) and target sequence only.

**Supplementary Table 4:** The sequences for SARS-CoV2 used to identify conserved regions to design crRNA guides is available as a separated Excel file

| **Endogenous genes** | |
|---|---|
| PPIB | Hs00168719_m1 (Thermo Fisher Scientific) |
| CXCR4 | Hs00607978_s1 (Thermo Fisher Scientific) |
| KRAS | Hs00364284_g1 (Thermo Fisher Scientific) |
| GAPDH | Hs02786624_g1 (Thermo Fisher Scientific) |
| **PB1 WSN/33** | |
| Forward | ATCTTTGAGACCTCGTGTCTTG |
| Reverse | CAGCAGGCTGGTTCCTATTTA |
| Probe | ACACGAGTGGACAAGCTGACACAA |
| **PB2 WSN/33** | |
| Forward | GTCAGTGAAACACAGGGAACA |
| Reverse | CCAACACTGATTCAGGACCATTA |
| Probe | ACTTACTCATCGTCAATGATGTGGGAGA |

**Supplementary Table 5:** Primer and probes (5' to 3') used for qPCR assays.
